# Comparative assessment of allelopathic herbicidal actions and total phenolics content of seven Indian medicinal plants

**DOI:** 10.1101/2020.12.23.423709

**Authors:** Souren Goswami, Sanjib Ray

## Abstract

Allelopathy is a vital ecological process that influences the dynamics of ecological succession, structure, and composition of plant communities. The present study aimed to evaluate a comparative account of allelopathic inhibitory actions in relation to total phenolic contents of aqueous extracts from seven traditionally used medicinal plants of India. *Triticum aestivum* and *Cicer arietinum* seedlings were used to test seed germination inhibition, seedling growth retardation and branch root sprouting inhibition. *Allium cepa* root tip cells were used for mitotic index inhibition and abnormal cell percentage analysis. The correlation between the total phenolics content and allelopathic activities was analyzed. The different extracts show the varied degree of allelopathic inhibitory activity. Out of these seven used extracts, *Crinum asiaticum* leaf extract (CaLAE) showed the highest allelopathic inhibitory action and it could reduce 94.3 and 79.59% root and shoot growth respectively at 96 h of treatment (1 mg/mL) and that was increased to 96.18 and 93.78% respectively with 2 mg/mL. The quantitative phytochemical analysis also revealed that CaLAE also possess relatively higher amounts of total phenolics. The growth retardation effects of the extract are in accordance with the mito-depression and increased chromosomal abnormality in *A. cepa* root tip cells. In conclusion, *Crinum asiaticum* may be considered as a prospective source of allelochemicals for plant growth regulation and a source of commercial herbicidal products.

## 1. Introduction

Allelopathy is a vital ecological process that influences the dynamics of ecological succession, structure, and composition of plant communities (Weston, 1999). Allelopathy is defined as any direct or indirect, inhibitory, or stimulatory influence of plants on other plants due to the allelochemicals released into the environment (Putnam, 1988). The allelochemicals released from a plant into the environment may directly or indirectly modulate the neighboring other plants’ growth. Allelochemicals act by germination inhibition, causing damage to root-shoot, stunting their growth. These chemicals may be cytotoxic too as they play a vital role in the competition between plants and are also inhibitors of seed germination and natural herbicides (Inderjit et al., 2003; Abu-Romman et al., 2010). The physiological effects of allelochemicals include inhibition of photosynthesis and respiration, phytotoxic, cyto-genotoxic, nuclear condensation, and the influence on enzymatic activities (Putnam, 1988; Chaudhuri and Ray, 2015). The pharmaceutical and agrochemical industries widely use the active allelochemical components for the production of drugs, herbicides, and pesticides (Awodun and Ojeniyi, 1999; Parekh et al., 2005). These phytochemicals play a significant role in agro-ecosystems and alter the quality and quantity of crop products (Putnam, 1988).

The allelopathic actions are due to a variety of specific interactions of a single compound or mixtures at the cellular and molecular level (Chaudhuri et al., 2015). A number of the allelochemicals demonstrate biological activity and have been used in the agrochemical and pharmaceutical industries (Hamburger and Hostettmann, 1991). The allelochemicals such as saponins, alkaloids, terpenoids, phenolics, etc., are released by one plant species against herbivores, pathogenic microbes, or other competing plant species. The phenolics are the most abundant substance of phytochemicals that affect cell division and seedling growth (Rudner and Murray, 1996). The plant phenolics contribute to physiological functions such as seed maturation and dormancy and also play a vital role in the defence against predators and pathogens (Winkel-Shirley, 2002).

The different allelochemicals isolated from several plants and soils are of many chemical classes that may offer clues to new herbicide chemistry. Cinmethylin is an example of a commercial herbicide containing a natural product moiety. Cineole, a terpenoid, abundant in certain xerophytes, is the active portion of the molecule. If it is economically favorable to extract the active/precursor molecule from natural sources rather than synthesis, herbicide production would greatly be revolutionized. Many investigators have already attempted to exploit allelopathy as a weed control strategy with promising results (Lehle and Putnam, 1983; Putnam and DeFrank, 1983; Rizvi et al., 1987). The water extracts of sunflower (*H. annuus*) and sorghum (*S. bicolour*) were reported as natural bio-herbicides which can be effectively used in plant protection (Jamil et al., 2009; Dayan et al., 2009; Macias et al., 2010; Grisi et al., 2012). Recently we have investigated comparative antioxidant potentials and total phenolics of these seven plants. Moreover, phytochemicals profile, traditional and pharmacological applications of the plant products were reviewed in detail (Goswami and Ray, 2017).

In continuation of the earlier work, here we aimed to assess a comparative account of allelopathic seedling growth inhibitory actions of seven plants extracts and to correlate with their total phenolics contents. For the laboratory-based allelopathic inhibitory activity assessment, seed germination, seedling growth, and branch root sprouting inhibitions were studied in wheat (*Triticum aestivum*) and chickpea (*Cicer arietinum*). The mitotic index inhibition and abnormal cell percentage were scored from the squashed root tip of *Allium cepa*. The novelty of the present study is that a comparative assessment of allelopathic potentials of these traditionally used antitumor plant products is done for the first time.

## 2. Materials and methods

### 2.1. Chemicals

Glacial acetic acid, Folin-Ciocalteu, sodium citrate, and methanol were obtained from BDH Chemicals Ltd, UK. Tannic acid powder was obtained from HIMEDIA Laboratories Pvt. Ltd., India. Ammonium molybdate and sulphuric acid were obtained from Qualigens. Aluminum chloride and sodium phosphate were obtained from Merck Specialities Pvt. Ltd., India. Benzene and ethyl acetate were obtained from SRL, Pvt. Ltd., Mumbai, India. All other chemicals used in this study were of analytical grade.

### 2.2. Plants collection and storage

Fresh leaves of *Cordia dichotoma*, *Holoptelea integrifolia, Crinum asiaticum*; whole aerial parts of *Croton bonplandianum*, *Cayratia carnosa* and *Scadoxus multiflorus* were collected from the Burdwan University campus in September of 2014. The collected plant parts were washed in tap water; shade dried, ground into small pieces, and pulverized using an electric grinder (Philips Mixer Grinder HL1605, Kolkata, West Bengal, India). *Trigonella foenum-graecum* seeds were collected fresh from a local shop and used intact. Plant species were taxonomically identified by Prof. Ambarish Mukherjee, Taxonomist, The University of Burdwan. Fully dried leaf powders were then stored in a separate airtight glass container for upcoming.

### 2.3. Extracts preparation

The 50 g of dried leaf powder/seeds were extracted twice in total 1 L (500 mL X 2) of distilled water in two equal phases of 12 h, at 60°C, in a hot water bath; extracts were filtered using Whatman filter paper #1 (GE Healthcare UK Limited, Buckinghamshire, UK). Then the filtrate was concentrated at 50°C, in a vacuum hot air oven for more than 6 h and the final volume was measured (Parekh and Chanda, 2007). The extracts were coded as CdLAE, stands for *C. dichotoma* leaf aqueous extract; HiLAE, for *H. integrifolia* leaf aqueous extract; CaLAE, for *C. asiaticum* leaf aqueous extract; CbAAE, for *C. bonplandianum* aerial parts’ aqueous extract; CcAAE, for *C. carnosa* aerial parts’ aqueous extract; SmAAE, for *S. multiflorus* aerial parts’ aqueous extract and lastly TfSAE; for *T. foenum-graecum* seed aqueous extract. The extracts were stored in −20°C freezer for future use. To determine the extract value and concentration, 25 mL (5 mL X 5 replicas) of the extract was evaporated to complete dryness in a hot (60°C) air oven.

### 2.4. Experimental plants models

Wheat (*Triticum aestivum*) seeds were used as a model to test germination inhibitory effects of the extracts. Wheat and chickpea (*Cicer arietinum*) seedling’s root and shoot growth inhibition was analyzed for phytotoxic and allelopathic activity assessments. Onion (*Allium cepa*) root tip cells were analyzed for mitotic index and cytotoxic index in terms of abnormal mitotic cells.

### 2.5. Wheat seeds germination inhibition

Healthy wheat seeds were surface sterilized by wringing in 1% sodium hypochlorite solution and allowed to germinate in a BOD under controlled conditions (25±2°C) on sterilized wet filter paper in autoclaved glass Petri dishes, containing the seven different test extracts in 1 mg/mL concentration of each. Similarly, seeds were maintained in Petri dishes, containing only distilled water was considered as untreated controls. 35 seeds in each Petridis and 5 replicas (total 175 seeds) of each test sample were maintained for 7 days. Treatment was given only once at the very beginning and every 24 h, the required amount of distilled water was added and mixed properly to maintain the moist condition.

### 2.6. Seedling growth inhibition

#### 2.6.1. Wheat

Healthy surface-sterilized wheat seeds were allowed for sprouting in distilled water for 36 h and then 15 seeds (approximately similar sprouting level) in each Petri dish with three replicas were taken for each group of test extracts. The group maintained using distilled water considered as the control. The dose of the treatment was 1 and 2 mg/mL. Root lengths were measured every 24 h of treatment up to 96 h.

#### 2.6.2. Chickpea

The similar size chickpea seeds were also surface sterilized and allowed for sprouting in distilled water for 48 h and then 15 seeds in each Petri dish with three replicas were taken for each group (1 and 2 mg/mL) of test extracts. The group maintained in distilled water considered as the control. Root lengths were measured in every 24 h of treatment up to 96 h. In chickpea seedlings, seminal root sprouting frequency was also measured at the final hour.

### 2.7. Mitotic index reduction and cytotoxic effects on onion root tip cells

For the evaluation of antiproliferative and cytotoxic effects of the seven plant extracts, the similar-sized onion bulbs were allowed to sprout roots in distilled water for 48 h at room temperature (25±2°C) (Greice et al., 2008), and then the roots were treated with the different extracts (1 mg/mL) for 24 h. The untreated control set was maintained in distilled water. The root tips were fixed in aceto-methanol (1:3 v/v) for 24 h (Ray et al., 2012). Aceto-orcein stained, fixed root apical meristematic cells (Sharma and Sharma, 1999) were squashed on slides, randomly coded, and observed under a light microscope for the altered mitotic index (MI) and chromosomal aberration if any. At least five slides were studied in each group.

### 2.8. Total phenolics contents and correlation analysis with allelopathic actions

The total phenolics contents of the seven plant products were estimated (Goswami and Ray, 2017; Makkar et al, 1993) and the data were used here for correlation analysis with the allelopathic actions.

### 2.9. Scoring and statistical analysis

Wheat and chickpea seedling’s root and shoot length were recorded and the growth retardation percentages were calculated. Student’s t-test was performed to analyze the significant (*p* <0.05, *p* <0.01, *p* <0.001) difference between the control and treated groups for root and shoot length. 2×2 contingency χ2-test (d.f. = 1) was performed for Mitotic index inhibition, abnormal mitosis, and apoptosis.

## 3. Result and discussion

### 3.1. Wheat germination inhibition

To test relative allelopathic potentials of the seven plant products, the germinating wheat seeds were treated with their aqueous extracts and germination inhibition % were analyzed. In this study, hot water was used as a polar solvent for chemical extraction as water is an essential requirement for seed germination and the water-soluble phytochemicals are extracted. Earlier reported that the allelopathic phytotoxins can be extracted with water and as well as a variety of organic solvents like petroleum ether, chloroform, ethyl acetate, methanol, etc. (Chaudhuri and Ray, 2015).

The data indicate a wide range (1.85 - 68.72%) of germination inhibition effects of the extracts on wheat seeds. Except for *Cd*LAE, all the other six extracts show a significant (5 - 0.1 %) level of germination inhibition at 72 h. The maximum germination percentage was recorded in the control set of seeds and the minimum germination percentage was recorded in *Ca*LAE treated samples. The decease order of germination percentage was determined as control (93.15±0.7) > CdLAE (91.43±0.9) > CbAAE (90.29±0.7) > CcAAE (83.43±1.1) > HiLAE (81.14±1.1) > TfSAE (70.86±1.1) > SmAAE (66.29±1.1) > CaLAE (29.14±1.1) (Table.1). Here, the CaLAE could reduce 68.72 % of wheat seedling germination on average. The allelopathic seed germination inhibition was also observed with the different aromatic and medicinal plants’ aqueous extracts by several scientists (Raoof et al., 2012; Marinov-Serafimov, 2010).

**Table 1.**
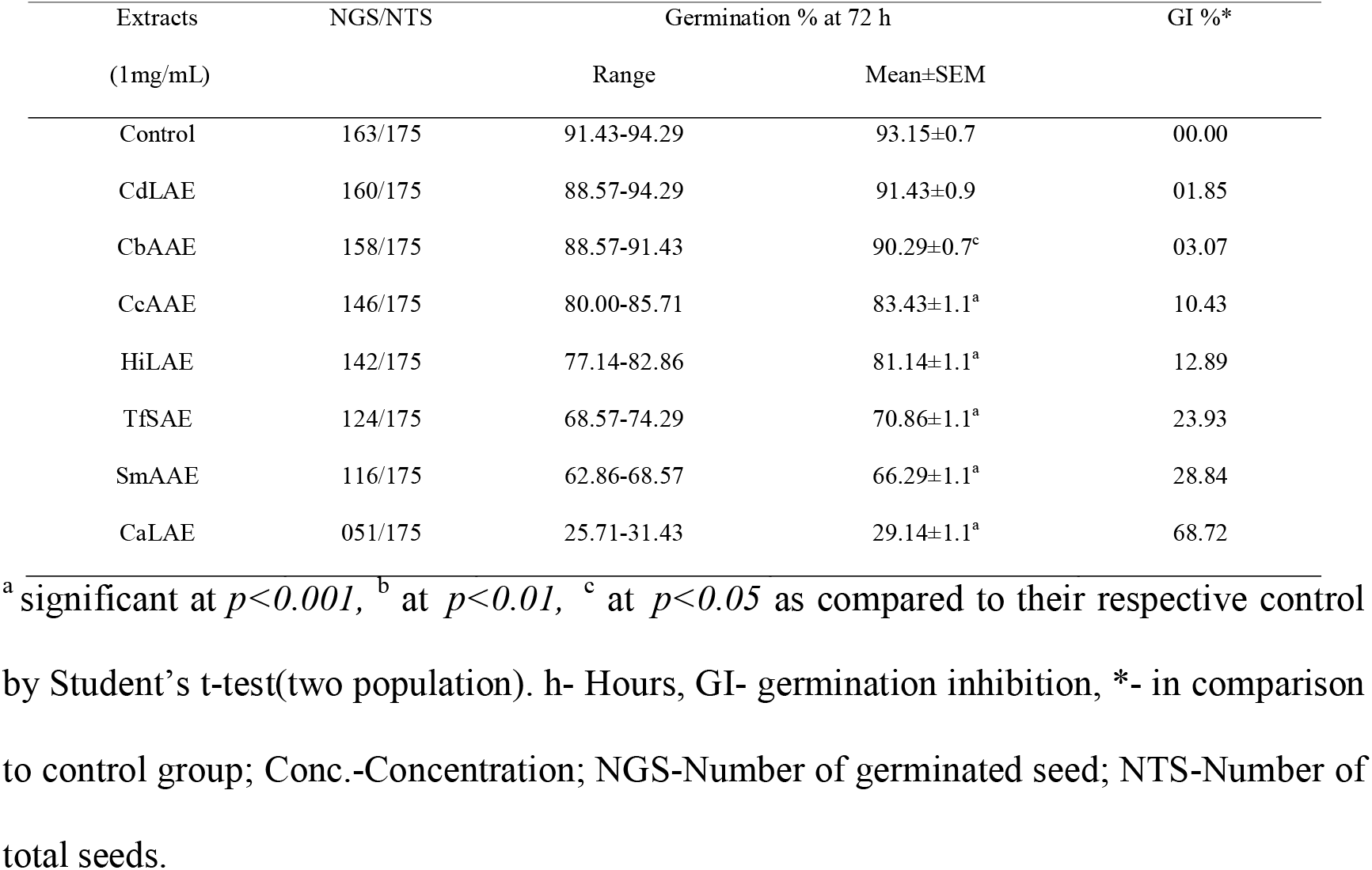
Influence of the different aqueous extracts on wheat seed germination.

### 3.2. Wheat seedling growth retardation

A comparative study was done among the aqueous extracts of seven plants using wheat seedlings to study their effect on root growth retardation. Seedlings are highly susceptible to allelochemicals due to high metabolic rate (Cruz-Ortega et al., 1998). A comparative study was done among the aqueous extracts of seven plants, using wheat seedlings to study their effects on root growth. Out of the seven extracts *Crinum asiaticum* leaf aqueous extract (CaLAE) was found to be the most effective growth retardant, followed by SmAAE, HiLAE, TfSAE, CdLAE, CcAAE. They all significantly retarded root growth at 0.1% level except CbAAE, which rather ameliorates it. Though CdLAE has no significant effect on wheat germination it reduced root-shoot growth significantly, whereas CbAAE is just the opposite in action. CaLAE has reduced on average 94.36 % root and 79.59 % shoot growth with respect to the control group at 96 h of treatment with the 1 mg/mL concentration, when the dose doubled it, reaches to 96.18 and 93.78 % respectively as compared to the untreated controls. The maximum root length (124.0±2.1 mm) was recorded from the experimental setup treated (1 mg/mL) with CbAAE and the minimum length (6.8±0.4 mm) was recorded from the CaLAE treated seeds. Similar to the roots, shoot growth was also affected by the extracts and was also in the same order (Table 2).

**Table 2.** Influence of the different aqueous extracts on wheat and chickpea seedlings root-shoot growth at 96 h of incubation.

| Extracts | Concentration (mg/mL) | Wheat (monocot) seedling |  |  |  | Chickpea (dicot) seedling |  |  |
| --- | --- | --- | --- | --- | --- | --- | --- | --- |
|  |  | RL (mm) | RGI % | SL (mm) | SGI % | RL(mm) Mean±SEM | RGI% | Branch root no. Mean±SEM |
|  |  | Mean±SEM |  | Mean±SEM |  |  |  |  |
| Control | 00 | 120.5±4.4 | 00.0 | 71.4±2.1 | 00.0 | 90.0±9.1 | 00.0 | 13.1±1.2 |
| CbAAE | 01 | 124.0±2.1 | -2.90 | 70.6±1.5 | 1.01 | 75.5±9.2 | 18.3 | 9.4±0.8<sup>c</sup> |
|  | 02 | 121.6±2.3 | -0.91 | 67.4±1.4 | 5.64 | 64.6±12.7 | 33.7 | 7.1±0.7<sup>a</sup> |
| CcAAE | 01 | 89.4±3.7<sup>a</sup> | 25.81 | 65.6±1.1<sup>c</sup> | 8.25 | 77.8±8.3 | 15.2 | 9.0±1.0<sup>c</sup> |
|  | 02 | 80.1±2.3<sup>a</sup> | 33.53 | 61.7±1.1<sup>a</sup> | 13.89 | 61.3±4.9<sup>c</sup> | 36.6 | 5.9±1.6<sup>b</sup> |
| CdLAE | 01 | 80.0±2.1<sup>a</sup> | 33.74 | 64.2±1.0<sup>b</sup> | 10.56 | 65.5±8.4 | 31.2 | 7.9±0.8<sup>b</sup> |
|  | 02 | 77.2±2.1<sup>a</sup> | 35.93 | 61.9±0.9<sup>a</sup> | 13.89 | 44.6±5.8<sup>a</sup> | 58.8 | 5.5±1.1<sup>a</sup> |
| TfSAE | 01 | 34.2±2.7<sup>a</sup> | 71.61 | 41.5±1.3<sup>a</sup> | 42.84 | 24.5±3.2<sup>a</sup> | 85.1 | 11.3±0.7 |
|  | 02 | 21.1±0.7<sup>a</sup> | 82.49 | 30.3±1.1<sup>a</sup> | 59.33 | 21.9±1.9<sup>a</sup> | 89.1 | 10.6±0.8 |
| HiLAE | 01 | 19.4±0.9<sup>a</sup> | 83.90 | 38.4±0.9<sup>a</sup> | 48.19 | 27.9±7.7<sup>a</sup> | 80.7 | 7.5±0.6<sup>a</sup> |
|  | 02 | 14.8±0.7<sup>a</sup> | 87.72 | 24.7±1.1<sup>a</sup> | 67.44 | 30.1±6.4<sup>a</sup> | 78.1 | 8.5±0.7<sup>b</sup> |
| SmAAE | 01 | 20.1±1.4<sup>a</sup> | 83.32 | 32.7±0.9<sup>a</sup> | 56.44 | 18.9±0.7<sup>a</sup> | 92.5 | 2.0±0.9<sup>a</sup> |
|  | 02 | 6.4±0.4<sup>a</sup> | 94.69 | 23.4±1.1<sup>a</sup> | 69.89 | 18.0±1.3<sup>a</sup> | 94.3 | 1.6±1.0<sup>a</sup> |
| CaLAE | 01 | 6.8±0.4<sup>a</sup> | 94.36 | 16.5±0.8<sup>a</sup> | 79.59 | 12.6±1.8<sup>a</sup> | 100.5 | 0.0±0.0<sup>a</sup> |
|  | 02 | 4.6±0.5<sup>a</sup> | 96.18 | 6.9±0.8<sup>a</sup> | 93.78 | 8.6±1.9<sup>a</sup> | 105.5 | 0.0±0.0<sup>a</sup> |

<sup>a</sup> significant at $p<0.001$ , <sup>b</sup> at $p<0.01$ , <sup>c</sup> at $p<0.05$ as compared to their respective control by Student's t-test (two population); h- hours; RGI- root growth inhibition in comparison to control group; SGI- shoot growth inhibition in comparison to control group; Conc- concentration; RL- root length; SL- shoot length.

Moreover, the present data indicate that roots are relatively more sensitive to the test extracts than shoots and it is in accordance with our previous report (Ray et al., 2012), which may because of root’s direct exposure to the phytochemicals rather shoots got them upon absorption. Allelopathic effects of plant extracts are well documented in the literature in terms of seedling growth inhibition (Grisi et al., 2012). The different aromatic and medicinal plants’ aqueous extracts have shown allelopathic seedling growth retardation effects (Raoof et al., 2012; Marinov-Serafimov, 2010). In the earlier studies with wheat seedlings, after treatment with a crude aqueous extract of *Ampelocissus latifolia* and *Synedrella nodiflora,* anomalies like rotting, necrosis, and complete atrophy of root hairs have been recorded (Chaudhuri and Ray 2014; Ray et al., 2013). Plant-based benchtop models are useful for the rapid evaluation of biological parameters including the toxicity of toxicants (Grisi et al., 2012). A potential allelopathic phytochemical Narciclasine 1, an Amaryllidaceae alkaloid, was isolated from the bulb of *Narcissus tazetta* based on wheat seedling growth inhibition (Ceriotti, 1967), which actually acts by affecting auxin transport (Hu et al., 2015), is now considered as a potent antitumor and anti-inflammatory agent (Furst, 2016). Incubation in the ethanolic root extract of *Calotropis gigantea,* exerted retarding effect on root emergence and growth of onion bulbs is evident of the cytotoxic effect caused by cytotoxic agents of the plant (Ravi et al., 2011).

### 3.3. Chickpea seedling root growth retardation

The seed germination, seedling growth and morphological alteration are generally considered for determining the allelopathic activity of plant extracts. The inhibitions of respiration, photosynthesis, and enzyme actions are the physiological effects of allelochemicals (Rice, 1984). Here, the HiLAE, TfSAE, SmAAE, and CaLAE could induce dose-dependent growth retardation on chickpea roots. Among them *C. asiaticum* leaf aqueous extract (CaLAE) was found to be the most effective, followed by SmAAE, TfSAE, HiLAE in both 1 and 2 mg/mL concentrations (Table 2). CdLAE, and CcAAE affected only in the higher (2 mg/mL) concentration, but CbAAE showed no effect. The maximum root length, 90±9.09 mm, was recorded from the untreated groups of the pea, while the minimum length, 8.63±1.93 mm, was recorded from the highest concentration, 2 mg/mL, of CaLAE at 96 h (Table 2). But if we look at branch root emergence, every test samples found to be effective except for TfSAE with zero effectiveness, whereas it reaches up to 100 % blockage in CaLAE in both the used concentrations.

*Euphorbia helioscopia* L. extracts show allelopathic actions on chickpea (*Cicer arietinum* L.), wheat (*Triticum aestivum* L.) and lentil (*Lens culinaris* Medic.) (Tanveer et al., 2010). The lettuce seedling growth was also significantly inhibited by Anthraquinones (Inove et al., 1992). If we consider at the cellular level, the root and shoot growth retardation is the outcome of the blockage or suppression in the cell division and induction of chromosomal aberrations and there exist positive correlations among them (Fiskesjo, 1985; Siddiqui, 2007; Usman et al., 2014).

### 3.4. Mitotic index reduction and cytotoxic effects on onion root tip cells

After counting of more than 4600 cells from each of the seven groups, patterns of MI alteration clearly indicate the differential mitotic inhibition effects of the different extracts on onion root tip cells, as there were significant (p<0.001) differences in mitotic index (MI) between the control and treated root tip cells except for *Cb*AAE and *Cc*AAE (Figure 1). The highest level of MI (12.62±1.2 %) was observed in CcAAE, followed by control (11.95±0.5 %), and that gradually decreases up to ^1^/_10_^th^ times of control, as in CaLAE (1.22±0.1 %) and SmAAE (1.17±0.04 %) treated tissues. CdLAE, HiLAE also exerted antiproliferative effects, with divisional rate as respectively 7.53 and 6.97 %, but TfSAE was far ahead with 2.42 % MI. HiLAE, TfSAE, SmAAE, and CaLAE also induced mitotic anomalies in 0.1 % significance level, with the highest values of 22.09±1.7 % abnormal out of total mitotic phases in SmAAE, followed by CaLAE (17.36 ±1.1 %). Figure 2 shows the different anomalies observed in plant extract treated onion root apical meristem tissues. The use of *Allium cepa* root tip cells for rapid assessment of cytotoxicity, genotoxicity, and mutagenic activities were accepted as a standard method, nearly a century ago (Camparoto *et al*., 2002, El-Shahaby et al., 2003; Sehgal et al., 2006; Angayarkanni et al., 2007; Fachinetto et al., 2007). The growth retardation effects of the extracts on *T. aestivum* and *C. arietinum* seedlings are in accordance with the MI inhibition and the increased chromosomal abnormality in *A. cepa* root tip cells. Similar type of root growth inhibitory and mitotic index reducing effects of leaf aqueous extract of *Clerodendrum viscosum* was observed in onion and wheat root tip cells (Ray et al., 2012), indicating, the extract with anti-proliferative effect, causing delay in cell cycle kinetics (Ray et al., 2013).

**Fig.1.**
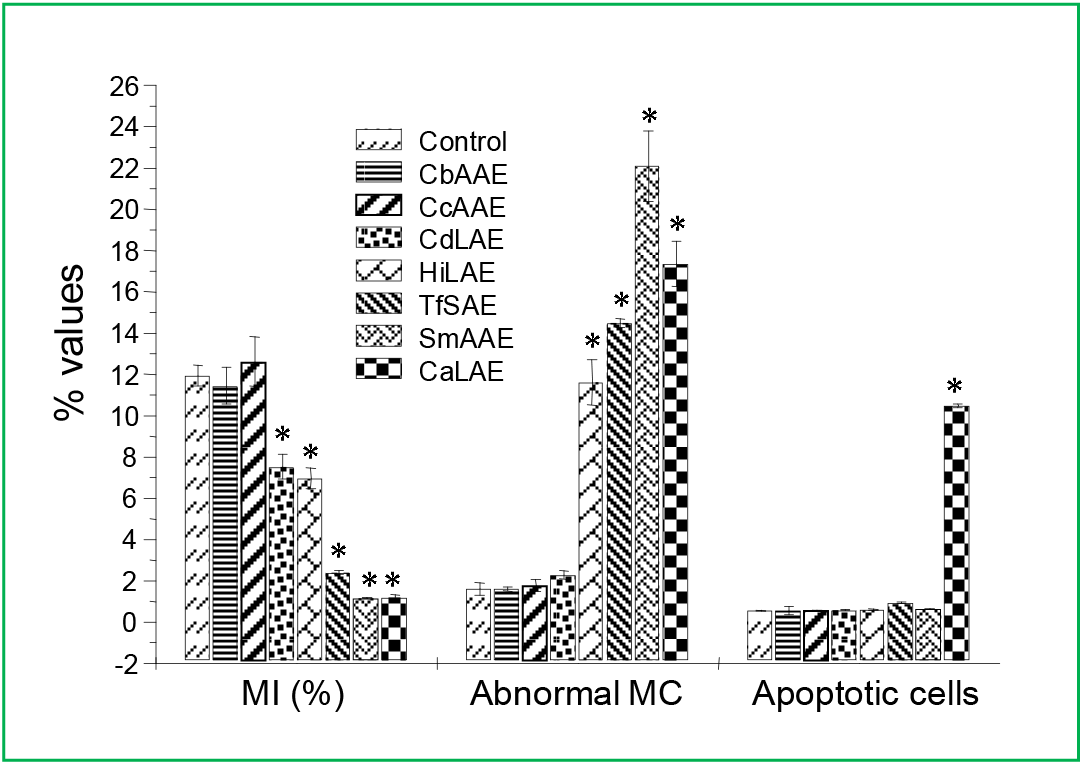
Effects of the different extracts on the mitotic index, abnormal mitosis, and apoptosis induction. *Significant at p<0.001 as compared to their respective control by 2-2 contingency χ2-test (d.f. = 1).

**Fig. 2.**
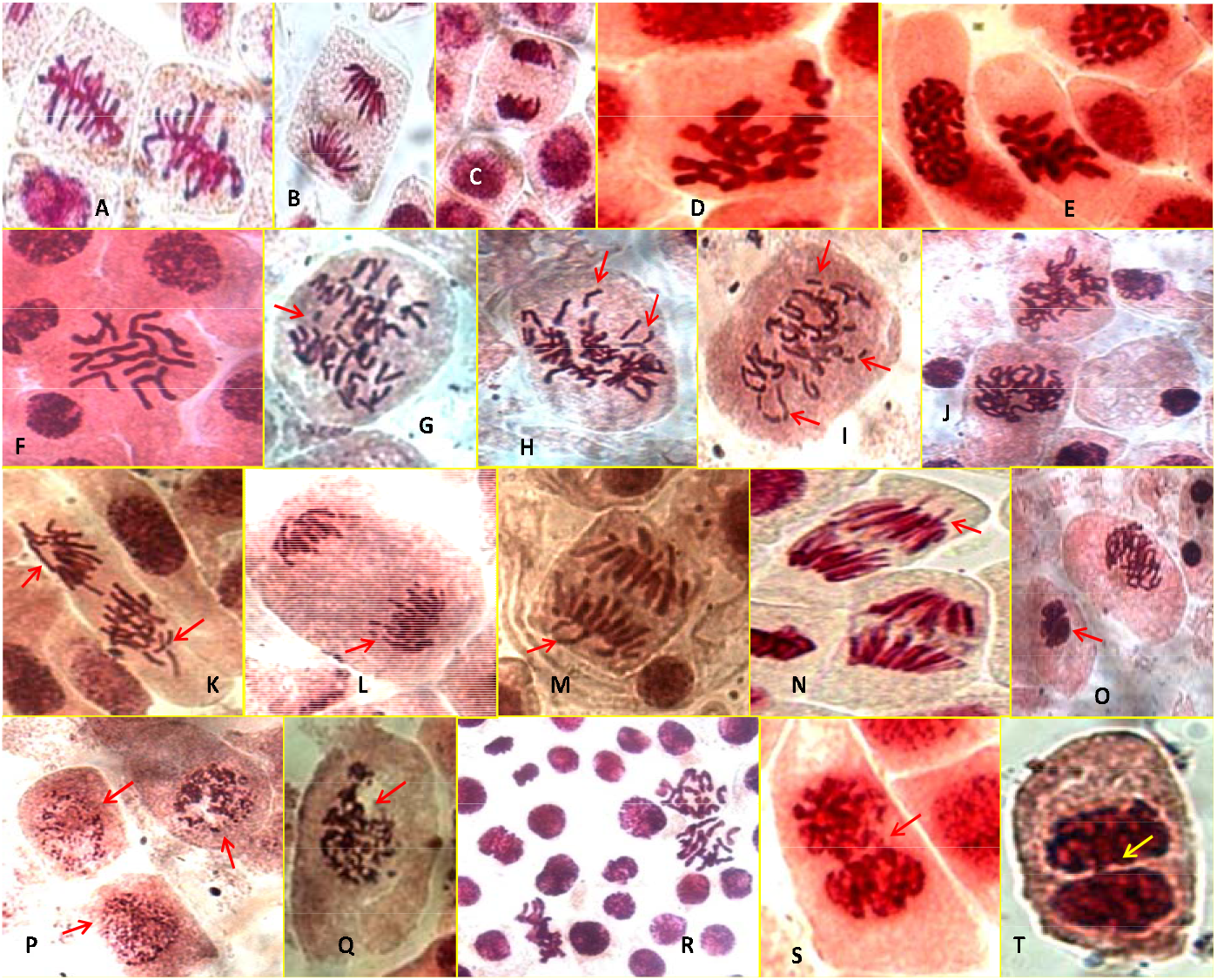
Photographs showing different types of abnormal mitotic phases in onion root apical meristem cells induced by different plant extracts (1 mg/mL), observed at 24 h of treatment; A, B, C- untreated divisional phases; D, E, F- c-metaphase; G, H, I- chromosomal breaks; J, R- sticky chromosome; K- vagrant chromosome; L- chromatin fragmentation; M, N- chromatin-bridge and vagrant chromosome; O- apoptotic nucleus and sticky chromosome; P, Q- fragmented apoptotic nucleus; R- sticky chromosome and c- metaphase; S, T- binucleate cells with a nuclear bridge;

In the present study, *Ca*LAE could show the highest allelopathic actions may be due to the presence of allelochemicals that could induce a reduction in mitotic index and apoptotic cell death. Different types of mitotic abnormalities induced in meristematic tissues of onion root by 1 mg/mL concentration of different plant extracts, at 24 h of treatment is shown in figure 2. Such abnormalities includes c-metaphase, chromosomal breaks, sticky chromosome, vagrant chromosome, chromatin fragmentation, chromatin-bridge, binucleate cells with nuclear bridge and also apoptotic cells with fragmented or blebed nucleus.

In this initial comparative study, SmAAE and CaLAE were found to be almost similar effective in antiproliferative action but the apoptosis-inducing ability was more in case of CaLAE and was the only significant at below 0.1 % level. Here, the CaLAE, representative of the Amaryllidaceae family is traditionally used to treat common cough-cold and throat disorder, vomiting, worm infestations, urinary disorders, bowel complaints such as colic, flatulence and even leprosy (Ghani, 1998; Walter, 1998; Zhanhe and Alan, 2000) and its reported antimicrobial and growth modulatory activities are in accordance with those properties (Ilavenil et al., 2010; Goswami et al., 2020).

### 3.5. Correlation between the total phenolics contents and allelopathic actions

The polyphenolics, plant secondary metabolites, are ubiquitously present in almost all plants. There exists a positive correlation between the reduced incidence of cancer and the consumption of phenolic-rich plant products (Hollman and Katan, 1999). Here, in figure 3 the correlation and regression analysis revealed a linear positive correlation between the total phenolic contents (data indicated in our previous publication, Goswami and Ray, 2017) and wheat germination inhibition (r = 0.816, r^2^ = 0.67), shoot growth inhibition (r = 0.862, r^2^= 0.74) and root growth inhibition (r = 0.892, r² = 0.795). Thus, the presence of total phenolics (Goswami and Ray, 2017) in the different extracts may have contributed directly to the allelopathic activities. Previous preliminary phytochemical analysis revealed, aqueous extract of *C. asiaticum* possess allelochemicals like alkaloids, terpenoids, tannins, and saponins (Goswami and Ray, 2017) and it is a promising source of antioxidants as well as antibiotics too (Goswami et al., 2020).

**Fig. 3.**
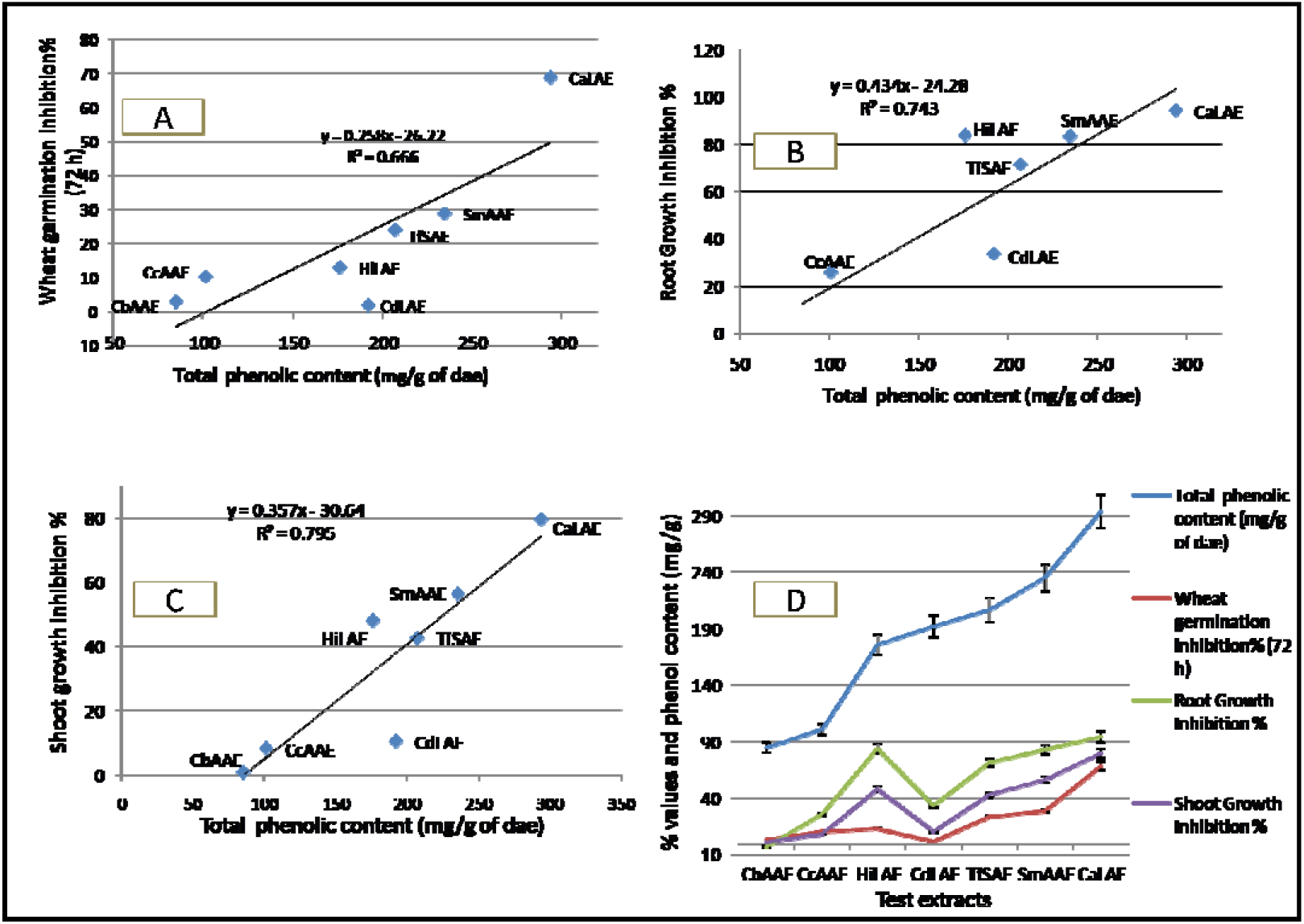
Showing a linear positive correlation between total phenolics contents of the seven different extracts and wheat germination inhibition % (A); wheat root growth inhibition % (B); wheat shoot growth inhibition % (C); in (D) wheat germination % and root-shoot growth inhibition % are plotted with their respective phenolics contents.

## 4. Conclusion

Globally thousands of researchers have identified numerous phytochemicals with growth retarding effects, on different models through simple table-top bioassays. Preliminary assessment of cell cycle modulation, mito-depression, and cytotoxicity are conveniently done through plant-based bioassays. They are considered simple and reliable methods and are widely used to detect toxicity. In this study, firstly we have screened the seven different plant aqueous extracts based on their MI modulation and induction of aberrations to these dividing cells. These ethnopharmacological approaches like screening for bioactivities, are very essential towards scientific validation of their traditionally claimed uses in herbal remedies and provide an indication in search of new active principles. The antiproliferative and cytotoxic bioactivities were considered as the primary criteria here to select a plant with a growth modulatory effect. The extracts of *C. asiaticum, S. multiflorus* and *T. foenum-graecum* showed significant dose-dependent root-shoot growth retardation effects in wheat. In chickpea seedlings, dose-dependent inhibition in seminal root emergence also found to follow the same pattern. After counting of more than 4600 cells from each of the seven groups, data indicate that the suppressing tendency of mitosis in CdLAE<HiLAE< TfSAE< SmAAE<CaLAE treated onion root tip cells as compared with the untreated group. *Crinum asiaticum* leaf extract was found to have the most phytotoxic and antiproliferative allelopathic actions among all the extracts tested. Thus, based on the above assessment of overall allelopathic inhibitory actions, phytotoxicity, and antiproliferation, *Crinum asiaticum* was found to be the most potent amongst the seven screened plant species and thus furthermore bioassays guided fractionation of this plant is on demand. In conclusion, *C. asiaticum* may be considered as a prospective source of commercial herbicidal products.

## Acknowledgment

The authors gratefully acknowledge the financial support of the UGC {F.No.42-563/2013 (SR) dt. 22.3.13}, UGC-DRS and infrastructural support of the Department of Zoology (DST-FIST and UGC-DRS Sponsored Department), The University of Burdwan, West Bengal, India.

## Abbreviations

CdLAE: *C. dichotoma* leaf aqueous extract
HiLAE: *H. integrifolia* leaf aqueous extract
CaLAE: *C. asiaticum* leaf aqueous extract
CbAAE: *C. bonplandianum* aerial parts’ aqueous extract
CcAAE: *C. carnosa* aerial parts’ aqueous extract
SmAAE: *S. multiflorus* aerial parts’ aqueous extract
TfSAE: *T. foenum-graecum* seed aqueous extract.

