## Supplementary tables for "Comparative assessment of allelopathic herbicidal actions and total phenolics content of seven Indian medicinal plants"

**Supplementary Data:**

**Table S1.** Showing results of correlation (R) and coefficient of correlation (R^2^) analysis amongst different variable factors

| **Correlation Between** | **The correlation of coefficient (R)** | **Coefficient of determination (R^2^)** | **Regression equation** |
| --- | --- | --- | --- |
| Total phenolics (mg/g of dae) in different extracts Vs Wheat germination inhibition % | 0.81609 | 0.666 | y = 0.258x - 26.22 |
| Total phenolics (mg/g of dae) in different extracts Vs Wheat root growth inhibition % | 0.862416 | 0.743 | y = 0.434x - 24.28 |
| Total phenolics (mg/g of dae) in different extracts Vs Wheat root growth inhibition % | 0.892129 | 0.795 | y = 0.357x - 30.64 |

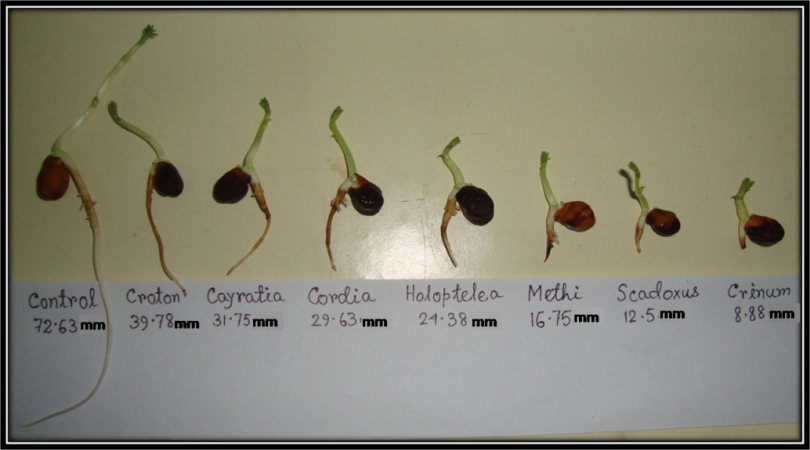

**Fig. S1.** *Chickpea* seedlings treated with 2 mg/mL concentration of different plant extracts, at 72 h. (* significant at *p<0.001* as compared to their respective control by Student’s t-test two population)

**Table S2.** Pooled data showing influence of the different extracts on mitotic index, abnormal mitotic phases and apoptosis.

| Dose  (1 mg/mL) | TC (%) | MI (%) | Mitotic phases | | | | Abn. MP % | Apop. cells % |
| --- | --- | --- | --- | --- | --- | --- | --- | --- |
|  |  |  | Pro (%) | Meta (%) | Ana (%) | Telo (%) |  |  |
| Control | 5308 | 11.95±0.5 | 47.55±1.5 | 15.99±2.7 | 19.32±1.4 | 17.15±1.9 | 1.61±0.3 | 0.55±0.01 |
| CbAAE | 4975 | 11.46 ±0.9 | 47.51±0.9 | 16.06±2.1 | 20.34±1.0 | 16.06±1.8 | 1.61±0.1 | 0.56±0.2 |
| CcAAE | 5030 | 12.62±1.2 | 47.64±1.1 | 16.80±1.3 | 19.24±0.9 | 16.32±1.1 | 1.78±0.3 | 0.56±0.01 |
| CdLAE | 4670 | 7.53±0.6^a^ | 45.15±1.9 | 17.29±0.4 | 16.19±0.5 | 21.38±1.0 | 2.29±0.2 | 0.60±0.02 |
| HiLAE | 5144 | 6.97±0.5^a^ | 42.65±1.2 | 17.37±0.5 | 19.59±0.6 | 20.39±1.9 | 11.62±1.1^a^ | 0.62±0.02 |
| TfSAE | 5103 | 2.42±0.1^a^ | 33.97±1.1 | 12.9±1.7 | 23.24±0.3 | 29.77±0.7 | 14.49±0.2^a^ | 0.94±0.04^c^ |
| SmAAE | 4677 | 1.17±0.04^a^ | 45.27±1.2 | 20.01±1.1 | 16.57±1.3 | 18.16±0.8 | 22.09±1.7^a^ | 0.64±0.02 |
| CaLAE | 5208 | 1.22±0.1^a^ | 63.99±1.1 | 12.53±0.9 | 9.66±1.2 | 13.81±1.1 | 17.36±1.1^a^ | 10.48±0.1^a^ |

^a^ Significant at *p*<*0.001*, ^b^ at *p<0.01* and ^c^ at *p<*0.05 as compared to their respective control with 2x2 contingency χ^2^-test with respective d.f. = 1. TC- total cells counted, MI-Mitotic Index (% values); Abn. MP- abnormal mitotic phases; Apop. Cells- Apoptotic Cells.

Total Phenolic content (mg/g of dae)

| Test samples | Total phenolic content (mg/g of dae) |
| --- | --- |
| Control | 0 |
| CbAAE | 85 |
| CcAAE | 101 |
| CdLAE | 192 |
| HiLAE | 176 |
| TfSAE | 207 |
| SmAAE | 235 |
| CaLAE | 294 |

**Fig. S2.** Total phenolic contents in the seven different plant extracts; estimated as tannic acid equivalent and expressed on per gm of dried extract mater basis.
